# R-loops shape chromatin architecture to promote balanced lineage allocation during differentiation

**DOI:** 10.1101/2025.06.23.661125

**Authors:** Chun-Hao Chao, Thomas G. Fazzio

## Abstract

R-loops—RNA:DNA hybrids that often form co-transcriptionally—are emerging as key regulators of genome function, yet their roles in shaping chromatin architecture and developmental potential remain incompletely defined. Here, we use inducible *Rnaseh1* expression in mouse embryonic stem cells (mESCs) to achieve acute, global R-loop depletion and systematically interrogate their impact on chromatin structure and lineage specification. We find that R-loop loss has minimal effect on steady-state gene expression or self-renewal. Instead, it leads to a striking reduction in H2A.Z occupancy at both active and bivalent promoters, accompanied by increased nucleosome density—revealing a previously unrecognized role for R-loops in maintaining promoter architecture. During gastruloid differentiation, R-loop-depleted mESCs exhibit a pronounced bias toward ectodermal fates, along with dysregulation of lineage-specific transcription factors and impaired cell-cell signaling. Consistent with these alterations, R-loop-depleted cells show widespread perturbations in gene regulatory networks across several early cell types. These findings uncover a critical role for R-loops in shaping the H2A.Z chromatin landscape and preserving balanced lineage trajectories during early development, offering new insights into the epigenomic regulation of stem cell fate.

## INTRODUCTION

Chromatin-associated RNAs are increasingly recognized as key regulators of nuclear processes, including gene expression, RNA processing, and epigenetic modification. These RNAs—such as long noncoding RNAs, enhancer RNAs, and upstream antisense RNAs—often act near their sites of transcription, modulating local chromatin states and transcriptional activity. While their importance in epigenetic regulation is well established, their biochemical properties pose challenges for functional characterization, and the mechanisms by which they influence chromatin architecture remain incompletely understood.

Among these RNAs, R-loops—typically co-transcriptional RNA:DNA hybrids that displace the non-template DNA strand—represent a major and dynamic class of chromatin-associated structures. Although unresolved R-loops are known to induce genome instability through replication-transcription conflicts and increased DNA fragility (Hamperl et al. 2017; Crossley et al. 2019; Hatchi et al. 2015), accumulating evidence suggests that R-loops also play regulatory roles in shaping the epigenome. For example, R-loops can inhibit DNA methylation by blocking DNA methyltransferases (Ginno et al. 2012) or recruiting demethylases (Arab et al. 2019; Sabino et al. 2022). In addition, R-loops promote the recruitment of the RNA methyltransferase METTL3-METTL14, leading to 6-methyladenosine (m6A) modification of RNA transcripts (Zhang et al. 2024), which impacts transcription termination, RNA stability, and translation (Yang et al. 2019; Ding et al. 2023). Finally, R-loops have been shown to co-localize with some active histone modifications, as expected given the requirement for transcription to generate R-loops.

We and others have previously shown that R-loops impact the binding of a few chromatin regulatory enzymes, including the Tip60-p400 histone acetyltransferase and histone exchange factor and the Polycomb Repressive Complex 2 (PRC2) (Chen et al. 2015; Alecki et al. 2020; Skourti-Stathaki et al. 2019). Notably, we previously observed a significant defect in the differentiation capacity of mouse embryonic stem cells (mESCs) that were partially depleted of R-loops (Chen et al. 2015). Despite these findings, a comprehensive understanding of how R-loops influence chromatin structure is lacking. Furthermore, the functional link between R-loops and mESC differentiation is not clear.

Here we comprehensively investigate the direct roles of R-loops in regulation of chromatin structure by generating an inducible *Rnaseh1* mESC model to test the acute effects of R-loop depletion on multiple features of chromatin architecture. Unlike chronic depletion models, this system enables examination of chromatin features without compensatory changes that may occur as cells adapt to *Rnaseh1* overexpression longer term. Upon acute reduction of R-loops genome-wide, most features of chromatin structure are unaffected, consistent with the modest phenotype of R-loop depletion in mESCs. However, we observe significant effects on two features of promoter-proximal nucleosomes: H2A.Z deposition and nucleosome occupancy. Interestingly, we observe R-loop-dependent H2A.Z incorporation not only at highly expressed genes, but also at developmental genes that are expressed at low levels in mESCs.

To better understand how these perturbations contribute to defects in mESC pluripotency, we performed gastruloid differentiation and single-cell RNA-seq. R-loop-depleted mESCs exhibit a skewed differentiation trajectory favoring ectodermal fates, accompanied by dysregulation of lineage-specific transcription factors, altered cell-cell signaling, and disruption of gene regulatory networks. Together, our findings reveal a previously unrecognized role for R-loops in shaping the H2A.Z chromatin landscape and maintaining balanced lineage specification, providing new insights into the epigenomic regulation of stem cell fate.

## RESULTS

### Acute R-loop Depletion Does Not Disrupt mESC Self-Renewal or Growth

To investigate the role of R-loops in shaping the epigenome in a system where they are functionally important (Chen et al. 2015), we generated a doxycycline (Dox)-inducible *Rnaseh1* (RHKI) mESC line. While *Rnaseh1* overexpression is a widely used method to deplete R-loops by degrading RNA within RNA:DNA hybrids, chronic overexpression can lead to cellular adaptation, potentially obscuring direct effects. To enable acute and uniform R-loop depletion, we engineered a system using the ZX-1 mESC line, which harbors a targeted rtTA transactivator at the *Rosa26* locus and a Cre-loxP cassette at *Hprt* (Iacovino et al. 2011). We replaced the Cre cassette with *Rnaseh1* via recombination (Supplemental Fig. S1A) and validated correct targeting and inducible expression. We determined *Rnaseh1* overexpression plateaued at 0.5 µg/ml Dox (Supplemental Fig. S1C) and remained stable for at least 96 hours post-induction (Fig. 1A). Finally, we observed that RHKI showed no significant difference in doubling time upon Dox addition (Supplemental Fig. S1D).

**Figure 1.**
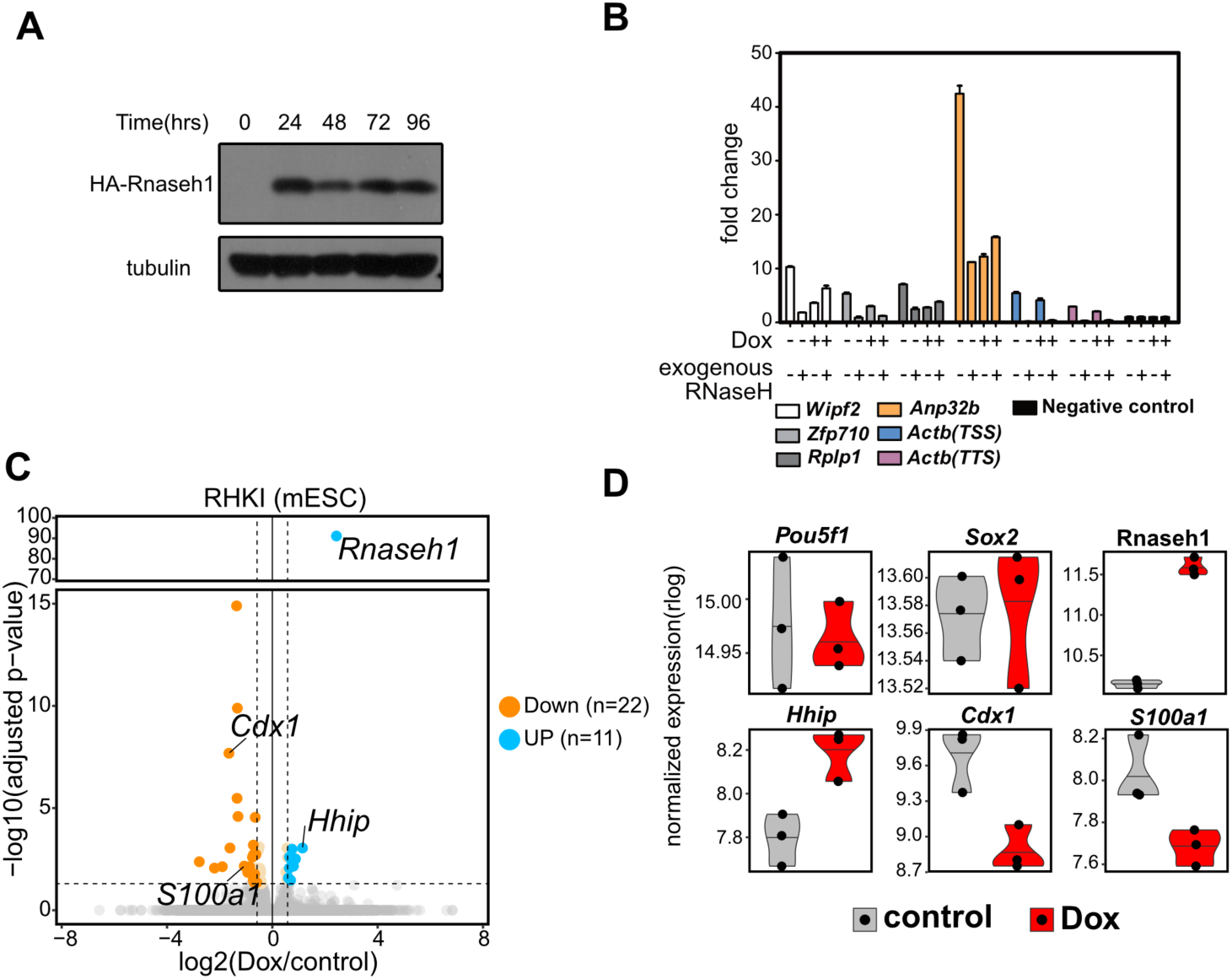
Acute R-loop disruption has minimal effects on mESCs. (A) Western blot of HA-RNaseH1 induction in response to Dox treatment (500 ng/ml) at 0, 24, 48, 72, 96 hours. Tubulin is included as a loading control. (B) Disruption of R-loops by acute induction of *Rnaseh1*. DRIP-qPCR is shown for several known R-loop locations: *Wipf2*, *Zfp710*, *Rplp1*, *Anp32b* and *Actb*, and a negative control locus after 48 hours of Dox treatment. Exogenous RNaseH treatment after DNA isolation is shown as an additional control. Mean and standard error of mean relative to the negative control locus are shown. (C) Changes in gene expression upon acute Rnaseh1 induction in undifferentiated mESCs. Shown is a volcano plot of significance relative to fold change in gene expression. 22 downregulated genes (orange) and 11 upregulated genes (blue) with the |log2FC| ≥ 0.58 and adjusted p-value ≤ 0.05 are indicated. (D) Gene expression of select genes from bulk RNA-seq. The expression level is transformed through regularized log transformation (rlog). *Pou5f1* and *Sox2* represent genes with no significant changes upon *Rnaseh1* induction and Rnaseh1, *Hhip*, *Cdx1* and *S100a1* represent genes significantly changed by *Rnaseh1* induction.

We next assessed R-loop depletion using DNA/RNA immunoprecipitation (DRIP)-qPCR across known R-loop-enriched loci (*Zfp710, Wipf2, Actb, Anp32b,* and *Rplp1*). Dox treatment led to a significant, though partial, reduction in R-loops, plateauing at ∼48 hours (Fig. 1B; Supplemental Fig. S1B). A control locus lacking R-loops showed minimal DRIP signal. These results are consistent with prior studies using constitutive *Rnaseh1* overexpression (Pires et al. 2023; Maul et al. 2017; Chen et al. 2015) and confirm that acute induction effectively reduces R-loop levels in mESCs.

To assess transcriptional consequences, we performed RNA-seq after 48 hours of Dox treatment. Only 33 genes were differentially expressed (fold change ≥ 1.5, adjusted *p* ≤ 0.05; Fig. 1C; Supplemental Table S1), with no enriched gene ontology categories. Pluripotency genes (e.g., *Pou5f1*, *Sox2*) were unaffected, though a few developmental regulators (*Cdx1*, *Hhip*) showed modest changes (Fig. 1D). These results indicate that acute R-loop depletion does not disrupt self-renewal or global gene expression but may prime cells for altered differentiation responses.

### R-loops Maintain Promoter-Proximal H2A.Z Enrichment in mESCs

R-loops have been associated with various epigenetic features, including DNA hypomethylation and histone modifications linked to active transcription, such as H3K4me3 and H3K27ac (Ginno et al. 2012; Sanz et al. 2016; Arab et al. 2019). However, it remains unclear whether R-loops directly regulate these features or simply co-localize with them due to shared association with transcriptional activity. To address this, we systematically profiled chromatin features following acute R-loop depletion using CUT&Tag and CUT&RUN.

We observed modest but reproducible changes in several histone marks, including H3K27ac, H3K36me3, and H3K27me3 (Fig. 2A; Supplemental Fig. S2A). Notably, the H3K27me3 response differed from that seen in chronic depletion models, highlighting the utility of acute perturbation. In contrast, we observed a striking and specific reduction in the histone variant H2A.Z occupancy following *Rnaseh1* induction (Fig. 2A), which was absent in Dox-treated parental ZX-1 cells (Supplemental Fig. S2C), confirming specificity. In mammals, H2A.Z is typically enriched at transcription start sites (TSS), including at both the −1 and +1 promoter-flanking nucleosomes, and at some enhancers (Ku et al. 2012; Raisner et al. 2005; Barski et al. 2007). Upon *Rnaseh1* induction, H2A.Z occupancy at TSSs was significantly reduced, while enrichment at distal DNase hypersensitive sites—used here as a proxy for enhancers—was only modestly affected (Supplemental Fig. S2B).

**Figure 2.**
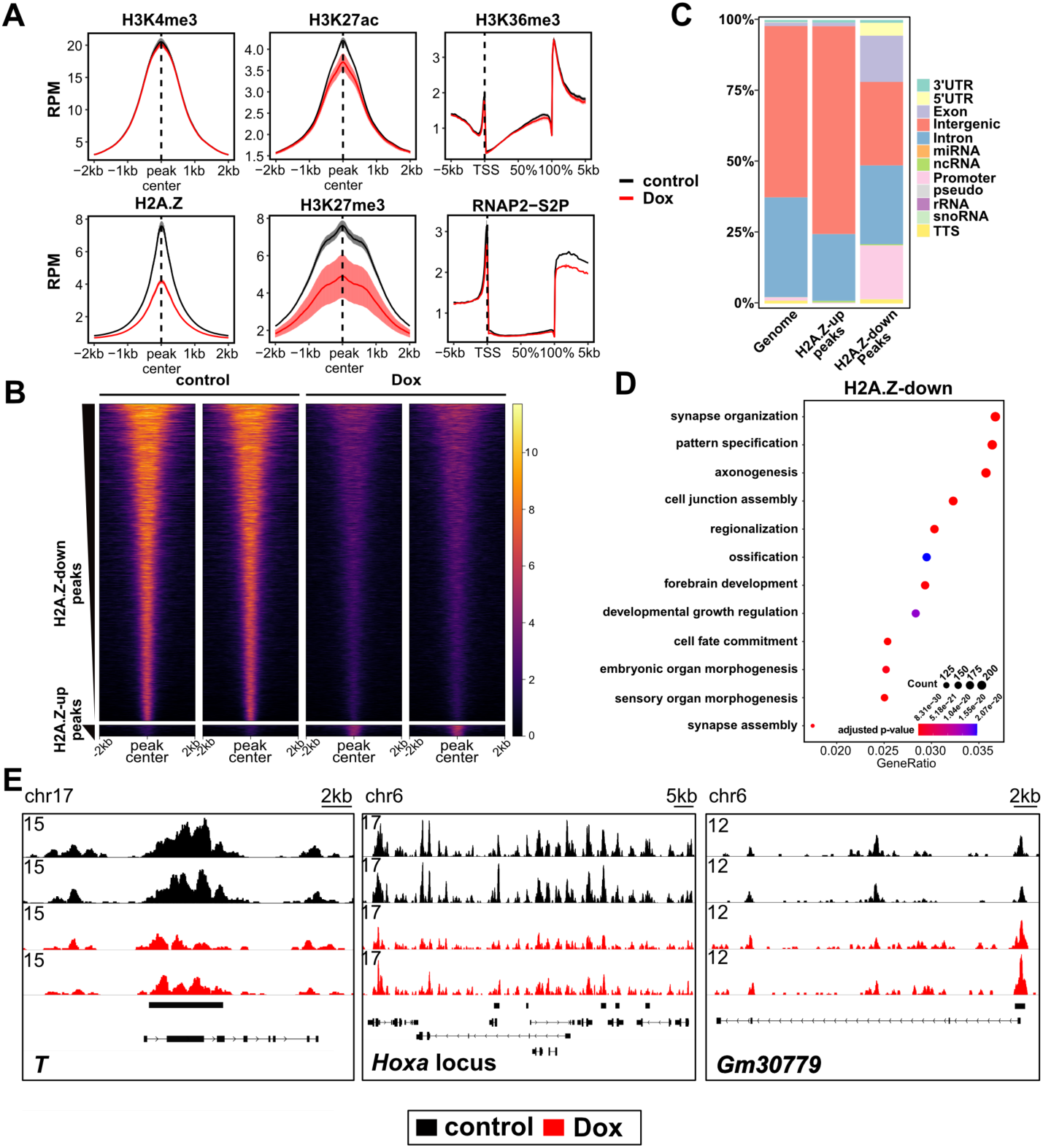
R-loop depletion causes genome-wide changes in H2A.Z occupancy in mESCs. (A) Enrichment of R-loop-associated epigenetic features. Aggregation of indicated features, measured by CUT&Tag surrounding peak centers from published ChIP-seq data (H3K4me3 and H3K27ac: GSE31039; H3K27me3: GSE123174; H2A.Z: GSE34483) or metagene profiles over gene bodies (H3K36me3 and RNAP2-S2P). Mean RPM (reads per 10 million) and standard error of mean are shown. (B) Heatmaps illustrating H2A.Z signal across peak centers of H2A.Z-down and H2A.Z-up peaks that differ between control and Dox treatment (defined with fold change ≥ 2 and FDR ≤ 0.05). Two representative replicates (out of four) of each condition are shown. (C) Quantification of genomic features of peaks belonging to the H2A.Z-up or H2A.Z-down peak set. The distribution of features across the mouse genome, denoted as ‘Genome’, is shown for reference. (D) Dotplot of GO category enrichment of the top 12 GO-terms of genes near H2A.Z-down peaks. A qvalue ≤ 0.05 was used as cut-off for GO category enrichment. (E) Genome browser tracks illustrating the effect of ectopic Rnaseh1 induction on H2A.Z enrichment. Black bars above gene models indicate significantly changed peaks. Two replicates each for control and Rnaseh1 induced (Dox) mESCs are shown.

Genome-wide analysis revealed 9,078 peaks with significantly reduced H2A.Z occupancy (H2A.Z-down) and only 319 with increased occupancy (H2A.Z-up) out of 61,420 total peaks (Fig. 2B). H2A.Z-down peaks were predominantly located at promoter-proximal regions, whereas H2A.Z-up peaks were more modest and enriched in intergenic regions (Fig. 2C), suggesting distinct roles for R-loops in regulating H2A.Z at promoters versus enhancers.

Gene ontology analysis of genes near H2A.Z-down peaks revealed enrichment for developmental and differentiation-related categories (Fig. 2D), despite low expression of these genes in self-renewing mESCs. Notably, H2A.Z loss was observed at promoters of key developmental regulators, including *T* and multiple *Hoxa* cluster genes (Fig. 2E). These findings suggest that R-loops facilitate H2A.Z deposition at both active and poised promoters, potentially priming them for activation during differentiation. Given the established role of H2A.Z in early development (Liu et al. 2022; Hu et al. 2013; Creyghton et al. 2008), its loss may compromise the fidelity of lineage commitment.

### R-loops Restrict Promoter-Proximal Nucleosome Occupancy

R-loops are frequently found near transcription start sites (TSSs) and transcription termination sites (TTSs). Thus, if R-loops promote nucleosome occupancy surrounding TSSs, their depletion could indirectly reduce H2A.Z enrichment at these sites. To test this, we first performed ATAC-seq in RHKI mESCs with and without acute *Rnaseh1* induction to assess changes in chromatin accessibility. Analysis of sub-nucleosomal fragments (≤120 bp), which mark nucleosome-free regions, revealed minimal differences in chromatin accessibility at TSSs and DNase I hypersensitive sites (Fig. 3A). No differential ATAC-seq peaks were detected genome-wide, and aggregate signal across all peaks remained unchanged (Supplemental Fig. S3B), suggesting that R-loop depletion does not broadly alter chromatin accessibility.

**Figure 3.**
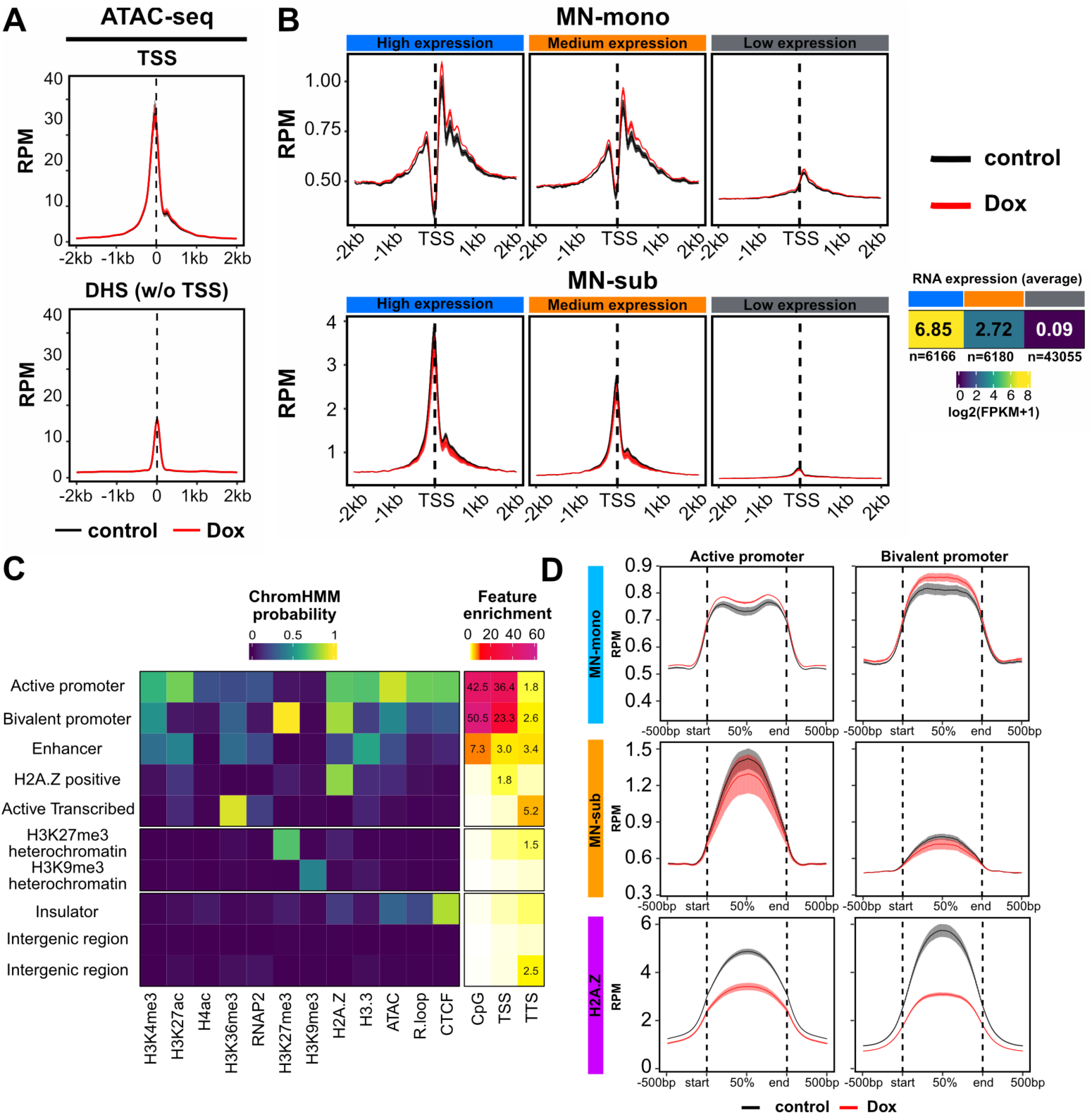
R-loop disruption increases nucleosome occupancy at active and bivalent genes. (A) Aggregation plot showing sub-nucleosomal ATAC-seq signal (1-120bp fragment size) across transcription start sites (TSSs) and TSS-distal DNaseI hypersensitive sites (DHS w/o TSS; DNase-seq: GSE37074) in control and Rnaseh1 overexpressing mESCs. Mean RPM and standard error of mean are shown. (B) Aggregation plot of MNase-seq signal in RPM surrounding the TSSs of high (n=6166), medium (n=6180) and low (n=43055) expression genes. Mean and standard error of mean are shown. MN-mono refers to mono-nucleosome sized fragments (150-200bp) and MN-sub refers to sub-nucleosome sized fragments (1-120bp). The three expression categories were determined using bulk-RNA-seq data from control mESCs and shown with the average of gene expression, log2(FPKM+1), in each category. (C) ChromHMM characterization of chromatin states from the epigenomic profiling data. Left: 10 states resulting from ChromHMM analysis of all ATAC-seq, CUT&Tag or CUT&RUN data measured in control mESCs. Right: enrichment of genomic features (CpG, TSS, and TTS) in each of the 10 states. (D) Aggregation plots of MN-mono, MN-sub and H2A.Z across active and bivalent promoter regions from ChromHMM. Mean RPM and standard error of mean are shown.

To more directly assess nucleosome positioning and occupancy, we performed MNase-seq, which captures nucleosome footprints across the genome (Lee et al. 2004; Song and Crawford 2010; Hughes and Rando 2014). We carefully titrated MNase digestion to ensure comparable digestion kinetics between conditions (Supplemental Fig. S3C) and analyzed both mononucleosome-sized fragments (MN-mono; 150–200 bp) and sub-nucleosomal fragments (MN-sub; ≤120 bp) at genes stratified by expression level. MN-sub fragments, which reflect nucleosome-depleted regions, were enriched upstream of TSSs and showed little change upon R-loop depletion (Fig. 3B), consistent with ATAC-seq results. In contrast, MN-mono fragments revealed a modest but reproducible increase in −1 and +1 nucleosome occupancy. These findings indicate that R-loops locally restrict nucleosome occupancy at promoter-proximal regions and rule out the possibility that reduced H2A.Z levels result from nucleosome loss.

To further explore the chromatin context of these changes, we trained a ChromHMM model using epigenomic data from control cells, identifying 10 distinct chromatin states (Fig. 3C). Upon *Rnaseh1* induction, we observed increased MN-mono signal specifically in states associated with active and bivalent promoters (Fig. 3D). Bivalent promoters—marked by both H3K4me3 and H3K27me3—are typically found at developmental genes that are modestly transcribed in mESCs but poised for activation. Notably, these are two of three states characterized by high probability of H2A.Z association and two of three states characterized by the presence of R-loops (Fig. 3C). In contrast, nucleosome occupancy remained largely unchanged across the remaining eight chromatin states (Supplemental Fig. S3D).

Together, these results demonstrate that R-loops help maintain a chromatin environment at promoter-proximal regions—particularly at active and poised developmental genes— characterized by H2A.Z enrichment and reduced nucleosome occupancy. While these changes do not disrupt self-renewal or steady-state gene expression, they may compromise the ability of mESCs to activate lineage-specific programs during differentiation.

### R-loop Disruption Alters Expression of Key Differentiation Regulators

Given the observed chromatin changes at developmental gene promoters, we next investigated how R-loop depletion affects mESC differentiation. Our previous work showed that chronic R-loop disruption impairs exit from pluripotency during embryoid body formation (Chen et al. 2015). To more comprehensively assess lineage specification, we employed the gastruloid model, in which mESC aggregates are exposed to defined stimuli to form 3D structures that mimic aspects of gastrulation (Beccari et al. 2018; Brink and Oudenaarden 2021).

Using RHKI mESCs, we initiated gastruloid differentiation following a two-day pre-treatment with or without Dox (Fig. 4A). Bulk RNA-seq was performed at 72 and 120 hours— representing mid and late stages of differentiation. Principal component analysis revealed that differentiation time was the primary driver of variance, with control and R-loop-depleted samples clustering by timepoint (Supplemental Fig. S1E), indicating that R-loop loss does not block differentiation per se. However, by 120 hours, we identified 245 upregulated and 228 downregulated genes in R-loop-depleted gastruloids (Fig. 4B; Supplemental Fig. S1F; Supplemental Table S1). Notably, several transcription factors critical for lineage specification were differentially expressed, including downregulation of *Cdx2* and *Foxa2*, and upregulation of *Neurog3* and *Sox5* (Fig. 4B–C). Trunk *Hox* genes (*Hox5–Hox11*), essential for posterior development (Brink and Oudenaarden 2021; Gouti et al. 2017; Ibarra-Soria et al. 2023), were also downregulated. Additionally, we observed altered expression of key signaling components, including *Fgfr4* (FGF pathway) and *Wnt5a*, *Wnt7a* (WNT pathway).

**Figure 4.**
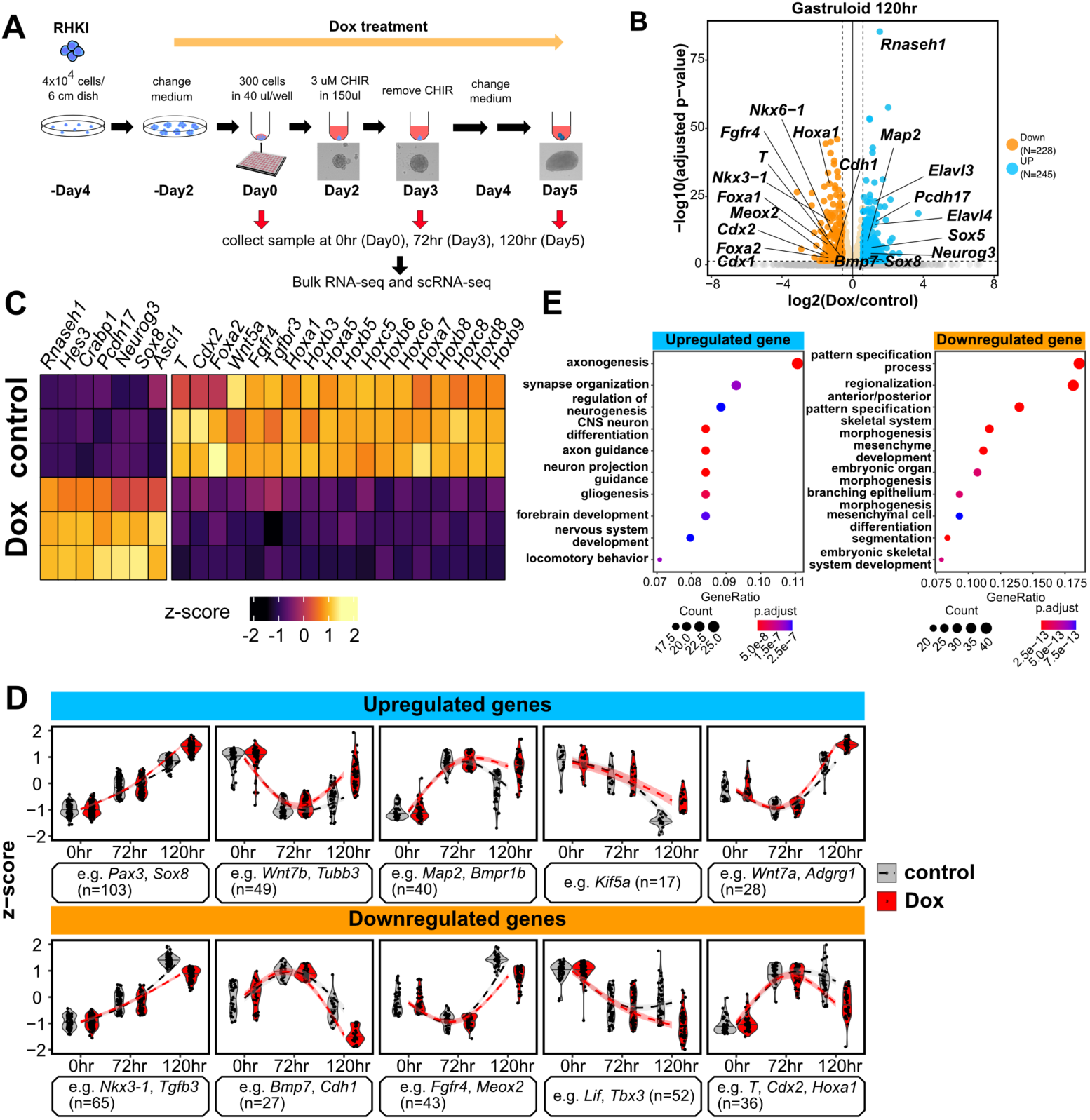
Key developmental regulators are dysregulated in gastruloids from R-loop depleted mESCs. (A) Schematic of gastruloid differentiation protocol and sample collection timepoints for bulk RNA-seq and scRNA-seq. RHKI was seeded at low density at Day −4 and Rnaseh1 was induced at Day −2 for two-days prior to differentiation. At Day 0, cells were re-plated for gastruloid differentiation (see Experimental Procedures). At Day 2, CHIR was added for one day to stimulate WNT signaling. (B) Volcano plot of bulk RNA-seq data at 120 hr of gastruloid differentiation, indicating changes in gene expression in Rnaseh1 overexpressing cells. Upregulated (n=228) and downregulated genes (n=245) are highlighted in blue and orange, respectively, with |log2FC| ≥ 0.58 and adjusted p-value ≤ 0.05 as cutoffs. (C) Example upregulated and downregulated genes in 120 hour gastruloids. Gene expression differences between control and *Rnaseh1* overexpressing cells are shown as z-scores. Three replicates for each condition were performed. (D) Violin plots of examples of upregulated and downregulated genes at 0, 72, and 120 hours with a regression line to indicate the expression trend across the time course. Expression trends were identified by k-mean clustering (see Experimental Procedures). (E) Dotplots of GO analysis results showing with top 10 GO-terms for upregulated and downregulated genes. GO categories shown are with qvalue ≤ 0.05.

To explore the dynamics of these changes, we classified differentially expressed genes based on their temporal expression trajectories (Fig. 4D). For example, *Pax3* and *Sox3*, which promote neural differentiation, were progressively induced and further elevated in R-loop-depleted cells. In contrast, other developmental regulators showed reduced expression in the absence of R-loops. Gene ontology analysis revealed that upregulated genes were enriched for neuronal differentiation, while downregulated genes were associated with pattern specification and somite development (Fig. 4E). These findings suggest that R-loop depletion may skew differentiation trajectories—enhancing ectodermal/neural lineages while suppressing mesodermal and endodermal lineages. However, bulk RNA-seq cannot identify the steps at which differentiation skewing may occur or which cell types are most affected.

### R-loop Disruption Skews Lineage Allocation During Gastruloid Differentiation

To directly assess whether R-loop depletion alters lineage specification, we performed single-cell RNA sequencing (scRNA-seq) during gastruloid differentiation in the presence or absence of *Rnaseh1* overexpression. This approach enabled high-resolution profiling of cell states and developmental trajectories.

Using the same experimental design as for bulk RNA-seq, we integrated our scRNA-seq data with two published gastruloid datasets (Suppinger et al. 2023; Rosen et al. 2022) for joint analysis. After applying a unified preprocessing pipeline (see Experimental Procedures), we observed strong concordance across datasets, with overlapping cell clusters in UMAP space (Supplemental Fig. S4A–B). Minor differences were attributable to sampling timepoints—for example, our 0-hour samples (undifferentiated mESCs) were unique, while 72- and 120-hour samples clustered closely with published data. To annotate cell types, we leveraged reference scRNA-seq data from mouse embryos (E3.5–E8.5) (Qiu et al. 2022; Cheng et al. 2019; Mohammed et al. 2017; Kearns et al. 2023; Pijuan-Sala et al. 2019) and used CellMatch to assign gastruloid clusters to embryonic lineages. We identified clusters corresponding to all three germ layers: ectoderm (e.g., Caudal Neuroectoderm, Neuromesodermal Progenitors, Spinal Cord, Neuron-like Cells), mesoderm (e.g., Primitive Streak, Paraxial Mesoderm, Splanchnic Mesoderm), and endoderm (e.g., Definitive Endoderm/Gut) (Fig. 5A–B). All clusters were present in both control and R-loop-depleted samples and expressed expected marker genes (Supplemental Fig. S4D), confirming their identities.

**Figure 5.**
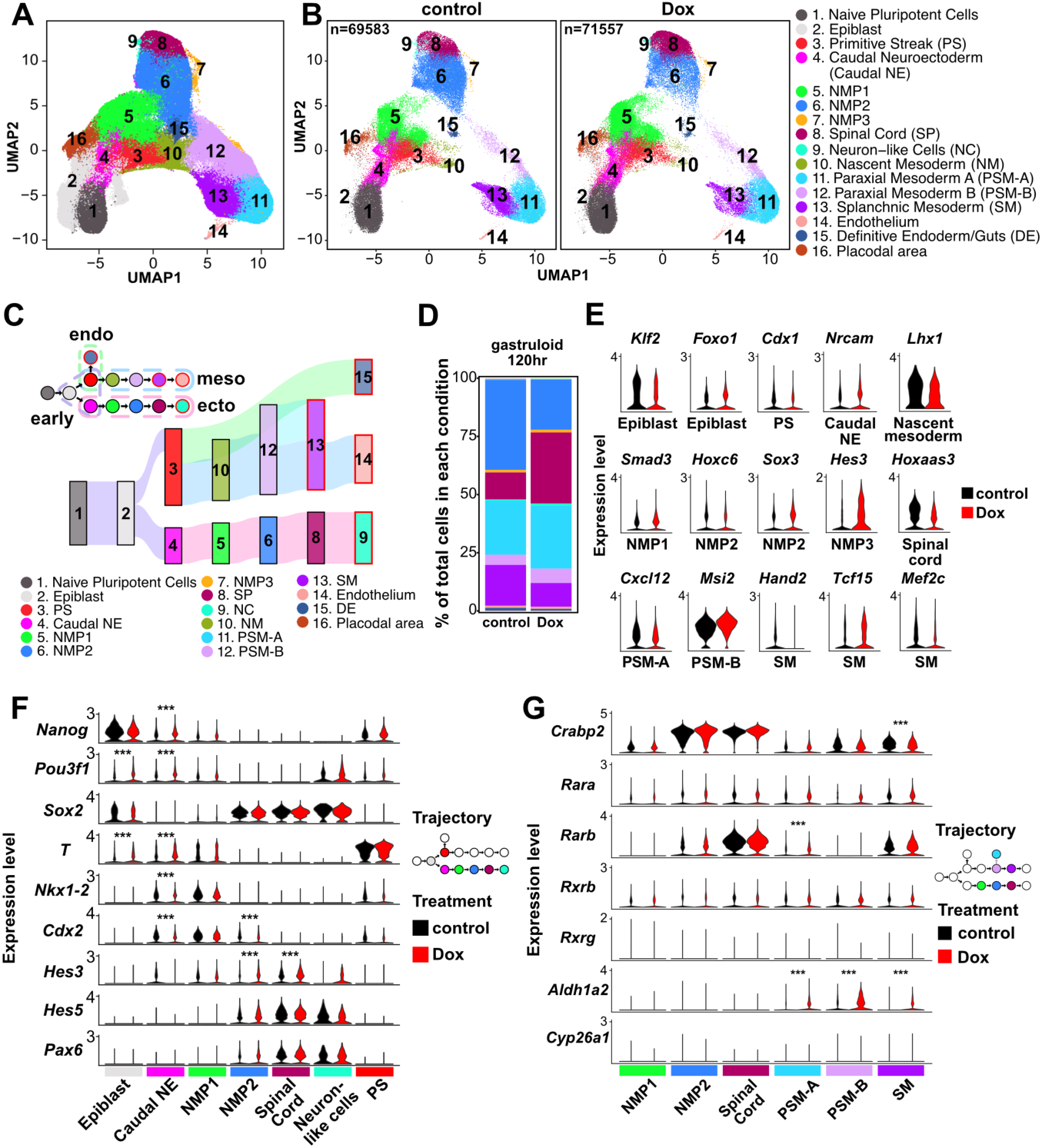
R-loop depletion skews differentiation in gastruloids. (A) Clusters of scRNA-seq data comprised of two published datasets combined with control and Dox-induced RHKI cells from this study, shown as a UMAP. Cell type annotations are shown in (B). NMP: Neuromesodermal Progenitors. (B) UMAP of scRNA-seq from RHKI across three timepoints, split by treatment condition: control (left) and Dox (center). Cell annotations are indicated to the right. (C) Below: Sankey plot showing the lineage trajectory of control scRNA-seq data. Above: Simplified lineage diagram with dotted lines indicating the cell types corresponding to each germ layer or early progenitor. (D) Proportions of total cells in control and Dox conditions at 120 hours of gastruloid differentiation. Colors correspond to cell types shown in (C). (E) Violin plots of differentially expressed genes (DEGs) between control and Dox conditions for different differentiated cell types. DEGs were called as described in the Experimental Procedures with adjusted p-values ≤ 0.05, |log2FC| ≥ 0.4 and the percentage of expressed cells ≥ 0.25. PS: Primitive Streak, Caudal NE: Caudal Neuroectoderm, NMP: Neuromesodermal Progenitors, PSM-A: Paraxial Mesoderm A, PSM-B: Paraxial Mesoderm B, SM: Splanchnic Mesoderm. (F) and (G) Violin plots of critical genes among cell types in ectoderm trajectory (F) and genes related to retinoic acid signaling among NMP/spinal cord and PSM/SM (G). *** indicates adjusted p-value < 0.0001.

To explore lineage dynamics, we performed trajectory inference by integrating our data with time-resolved gastruloid datasets and applying a modified pyVIA algorithm that incorporates both transcriptomic similarity and timepoint information. This analysis reconstructed a comprehensive differentiation map with three major germ layer trajectories and revealed key lineage decision points (Fig. 5C; Supplemental Fig. S5A). Comparing transition probabilities between conditions, we found that R-loop depletion increased transitions from Epiblast to Caudal Neuroectoderm and from Spinal Cord to Neuron-like cells (Supplemental Fig. S5B). These changes were reflected in altered lineage proportions: Neuromesodermal Progenitors (NMP1/NMP2) and spinal cord were overrepresented in R-loop-depleted samples at 72 and 120 hours, respectively (Fig. 5D; Supplemental Fig. S5C–D), suggesting accelerated and biased differentiation toward neural fates. These findings indicate that R-loop loss perturbs early lineage decisions, favoring ectodermal over mesodermal and endodermal outcomes.

### R-loop Depletion Alters Expression of Lineage-Specific Transcription Factors

To uncover the molecular basis of lineage skewing, we examined transcription factor (TF) expression across cell types. Many TFs critical for lineage specification were differentially expressed upon *Rnaseh1* induction (Table S2). For example, *Klf2* was downregulated in epiblast, *Cdx1* and *Eomes* in primitive streak, and *Mxi1* was upregulated in paraxial mesoderm A (Fig. 5E; Supplemental Fig. S5E). In splanchnic mesoderm, we observed reduced expression of *Hand2* and *Mef2c*, two TFs essential for cardiac development (Lin et al. 1997; Anderson et al. 2017; Liu et al. 2009; Miquerol and Kelly 2013), suggesting compromised mesodermal potential.

Given that lineage skewing appeared to originate at the epiblast stage, we closely examined TF expression at this transition point. *Pou3f1*, a neural-promoting TF expressed early in neurogenesis, was upregulated in both epiblast and caudal neuroectoderm in R-loop-depleted cells (Fig. 5F). Its early induction may contribute to the observed bias toward ectodermal differentiation. Neuromesodermal progenitors (NMPs), which co-express *Sox2* and *T*, can adopt neural or mesodermal fates depending on the relative expression of these factors (Chalamalasetty et al. 2011). Although we did not capture the double-positive NMP state, the trajectory analysis indicated a transition from NMP1 to NMP2, characterized by increased *Sox2* and decreased *T* expression, suggesting neuronal fate commitment of the NMPs. This transition was accompanied by upregulation of downstream neural TFs such as *Hes3*, *Hes5*, and *Pax6*, further supporting ectodermal commitment. We also observed downregulation of TFs involved in posterior axial patterning and early lineage specification, including *Nkx1-2*, *Cdx1*, and *Cdx2*. Interestingly, *T* was upregulated in caudal neuroectoderm, which may enhance NMP formation and contribute to altered lineage dynamics (Henrique et al. 2015).

Together, these findings reveal that R-loop depletion induces cell type–specific changes in key lineage-specific transcription factors, consistent with the observed shifts in lineage allocation. These results highlight a previously unrecognized role for R-loops in regulating transcriptional programs that guide early cell fate decisions.

### Elevated retinoic Acid Signaling Reinforces Lineage Skewing in R-loop-Depleted Gastruloids

During gastrulation, WNT signaling is essential for primitive streak formation and anterior-posterior patterning (Morgani and Hadjantonakis 2019). In gastruloids, treatment with the WNT agonist CHIR99021 is critical for elongation and spatial patterning of lineage-specific gene expression (Brink et al. 2014; Suppinger et al. 2023; Cermola et al. 2021), and alteration of growth factor exposure or simulation of cell–cell communication (CCC) can drive gastruloid differentiation toward specialized cell types (Brink and Oudenaarden 2021; Rossi et al. 2021, 2022).

Given the observed lineage skewing in R-loop-depleted gastruloids, we therefore investigated whether altered CCC contributes to this phenotype. Differential expression analysis revealed significant changes in genes involved in intercellular signaling. For example, *Fgfr1* (FGF/MAPK pathway) was dysregulated in multiple mesodermal populations, *Notch1* and *Notch2* were altered in paraxial mesoderm, and TFs such as *Smad3* and *Tcf7l1*, which mediate TGFβ and WNT signaling, were also affected (Fig. 5E; Supplemental Fig. S5F–G).

Notably, genes involved in retinoic acid (RA) signaling—a pathway critical for neuronal commitment of neuromesodermal progenitors (NMPs)—were significantly altered. *Aldh1a2*, the enzyme responsible for RA synthesis, was upregulated in mesodermal populations, while *Cyp26a1*, which degrades RA, remained low in both mesoderm and ectoderm (Fig. 5G). This imbalance suggests RA signaling from mesoderm to ectoderm may be enhanced in *Rnaseh1* overexpressing gastruloids. High expression of the RA adaptor *Crabp2* in NMP2 and spinal cord cells further supports increased RA signaling in ectodermal lineages. Slight upregulation of *Rarb* in mesoderm also suggests increased RA responsiveness, which may inhibit paraxial mesoderm differentiation (Russell et al. 2018).

To directly assess CCC changes, we used CellChat (Jin et al. 2025, 2021) to infer ligand–receptor interactions between cell types. This analysis revealed increased RA signaling from mesodermal populations (PSM-A, PSM-B) to ectodermal lineages (NMP2, spinal cord) in R-loop-depleted gastruloids (Fig. 6A). The most significantly enhanced interaction was *Aldh1a2–(Crabp2–Rarb)* (Fig. 6B; Supplemental Fig. S6A), suggesting a mesoderm-to-ectoderm RA signaling axis that reinforces neural fate commitment. Additional changes in FGF, Notch, and TGFβ signaling were also observed (Supplemental Fig. S6B). Together, these findings suggest that altered CCC—particularly increased RA-mediated signaling—contributes to the ectodermal bias observed in R-loop-depleted gastruloids.

**Figure 6.**
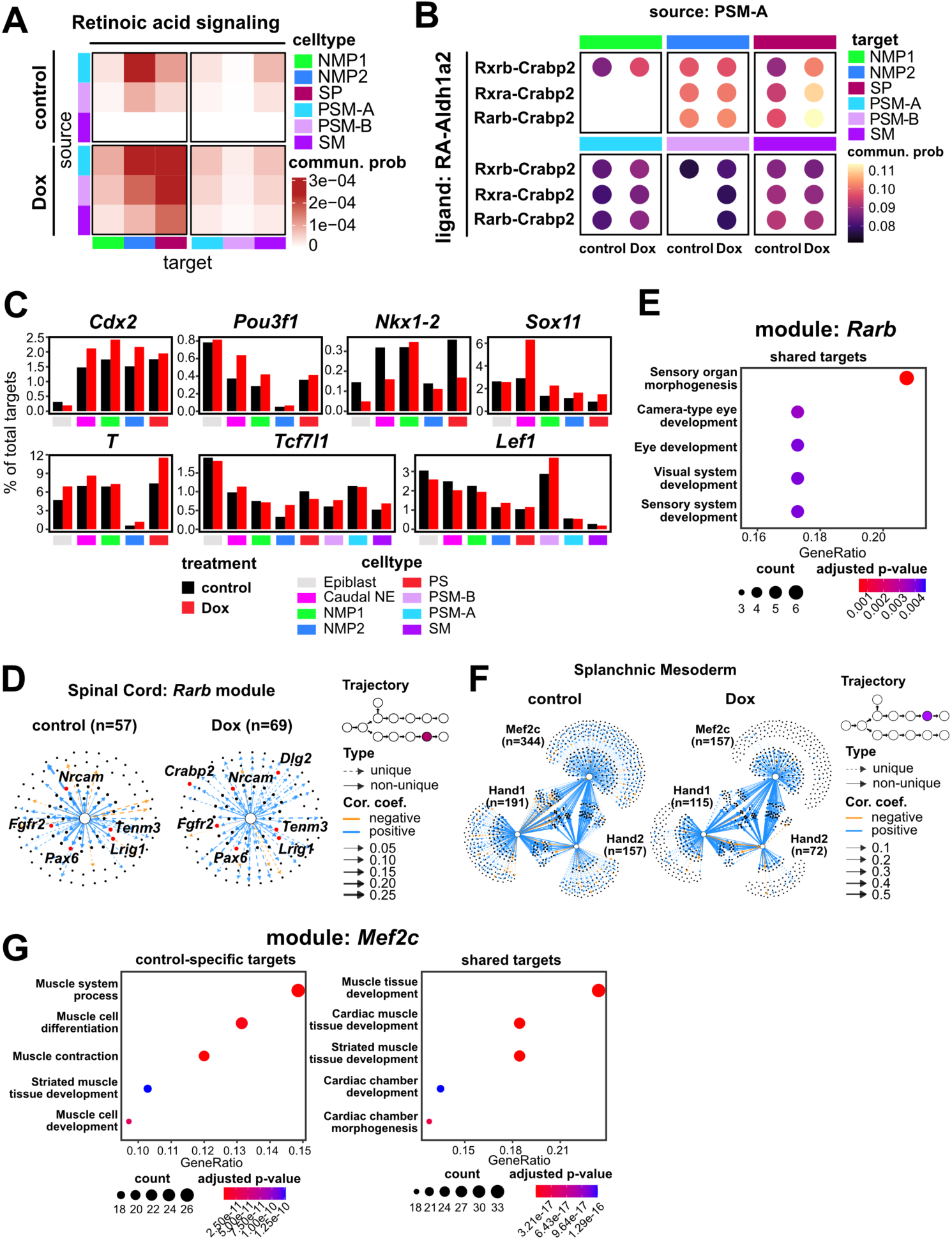
Alterations in signaling and lineage-specific GRNs in R-loop depleted gastruloids. (A) The heatmap of communication probability of retinoic acid signaling among mesoderm cells (PSM-A, PSM-B, SM) and ectoderm cells (NMP1, NMP2, spinal cord). The sources and targets of inferred RA signaling are indicated. (B) The communication probabilities of indicated ligand-receptor pairs in retinoic acid signaling from PSM-A to mesoderm cell types (PSM-A, PSM-B, SM) and ectodermal cell types (NMP1, NMP2, spinal cord). (C) Barplots depicting the proportion of targets in the indicated module related to the total target in each specific cell type/treatment combination. (D) Network plots illustrating TF-target relationships in the Rarb module for spinal cord. The trajectory diagram highlights the cell type in the trajectory map. ‘Unique’ edges indicate TF-target pairs that only exist in a specific combination of cell type and treatment, shown with a dashed line, whereas solid lines denote non-unique pairs. Edge thickness indicates the correlation strength of the TF-target pair. The line color of each edge indicates positive or negative correlation between TF and its targets. (E) Dotplot of GO analysis results for target genes in Rarb module. Shared targets indicate genes are positively correlated and shared between control and Dox. A qvalue ≤ 0.05 was used as cut-off for GO category enrichment. (F) Network plots illustrating TF-target relationships in the Hand1, Hand2 and Mef2c modules for splanchnic mesoderm. Labels are as in (D). (G) Dotplot of GO analysis results for target gene categories in Mef2c module. The positively correlated, control specific targets (left) and targets shared between control and Dox (right) are shown. A qvalue ≤ 0.05 was used as the cut-off for GO category enrichment.

### R-loop Depletion Rewires Gene Regulatory Networks to Promote Neuronal Differentiation

To understand how altered CCC and transcription factor activity contribute to lineage skewing, we reconstructed gene regulatory networks (GRNs) using scMTNI (Zhang et al. 2023), incorporating trajectory information to improve inference. Due to limited cell numbers in several terminal cell types, we focused on intermediate cell types.

We first examined GRN changes at the early bifurcation from epiblast to either caudal neuroectoderm or primitive streak. Upon *Rnaseh1* induction, we observed an increase in predicted *T* target genes across early lineages (Supplemental Fig. S6C), which remained after normalizing for total edges (Fig. 6C). We also observed expansion of *Sox11*, *Cdx2*, and *Pou3f1* regulatory modules in caudal neuroectoderm, alongside a reduction in the *Nkx1-2* module. Although WNT signaling is required for *T* induction, correlation analysis among *T*, *Lef1*, *Tcf7l1*, and *Nkx1-2* (Tamashiro et al. 2012) did not reveal major differences between conditions (Supplemental Fig. S6D), suggesting that altered WNT responsiveness is not the primary driver of *T* upregulation.

Consistent with enhanced RA signaling, the *Rarb* module was modestly expanded in spinal cord cells upon R-loop depletion (Fig. 6D). Shared *Rarb* targets were enriched for sensory system development, while Dox-specific targets included *Dlg2* and *Crabp2*, genes involved in neuronal differentiation and RA signaling, respectively (Sanders et al. 2022) (Fig. 6E; Supplemental Fig. S6E). These findings support a model in which RA signaling reinforces ectodermal fate commitment. In contrast, splanchnic mesoderm cells showed reduced expression of cardiogenic TFs *Hand2* and *Mef2c* (Fig. 5E), and GRN analysis revealed a marked reduction in *Hand1*, *Hand2*, and especially *Mef2c* target modules (Fig. 6F). GO analysis of *Mef2c* targets unique to control cells showed enrichment for muscle development, while shared targets were associated with cardiac development (Fig. 6G; Supplemental Fig. S6F), suggesting impaired cardiogenic potential in R-loop-depleted cells.

Together, these results demonstrate that R-loop depletion leads to widespread GRN rewiring, expanding neuronal gene programs while diminishing mesodermal and cardiogenic regulatory networks. These changes likely underlie the ectodermal bias and reduced mesodermal potential observed in R-loop-depleted gastruloids.

## DISCUSSION

In this study, we used an inducible *Rnaseh1* overexpression system to investigate the role of R-loops in chromatin regulation and mESC differentiation. Unlike previous studies that relied on long-term *Rnaseh1* overexpression—potentially confounded by cellular adaptation or transgene silencing—our system allowed us to examine the immediate consequences of R-loop depletion. Despite the broad potential of RNaseH1 to target RNA/DNA hybrids, acute R-loop depletion had minimal effects on steady-state mRNA levels or self-renewal, suggesting that the observed chromatin changes are direct consequences of R-loop loss. Notably, R-loop depletion was partial (∼2–3-fold reduction), consistent with prior studies (Maul et al. 2017; Chen et al. 2015), and may therefore underestimate the full impact of R-loops on chromatin architecture.

While most chromatin features—including histone modifications, RNA Polymerase II occupancy, and overall chromatin accessibility—were largely unaffected, we observed a pronounced reduction in H2A.Z occupancy at ∼20% of peaks and a modest increase in nucleosome occupancy. These changes were evident at both highly expressed genes and bivalent promoters, many of which regulate developmental programs. Given that H2A.Z is essential for proper differentiation (Hu et al. 2013), our findings suggest that R-loops are required to maintain H2A.Z deposition at key regulatory loci. This aligns with our previous observation that *Rnaseh1* overexpression reduces binding of Tip60-p400 (Chen et al. 2015), a chromatin remodeling complex involved in H2A.Z deposition. However, the mechanism linking R-loops to Tip60-p400 function remains unclear, as no physical interaction between RNA/DNA hybrids and any Tip60-p400 component was detected in a previous survey of RNA/DNA hybrid binding proteins (Wu et al. 2021b).

To explore the functional consequences of R-loop depletion, we used the gastruloid model to examine differentiation across all three germ layers. Bulk RNA-seq revealed dysregulation of key developmental genes, including *Cdx2* and multiple *Hox* genes—many of which also showed reduced H2A.Z occupancy in mESCs. Single-cell RNA-seq enabled reconstruction of differentiation trajectories that were consistent with prior gastruloid studies (Suppinger et al. 2023; Rosen et al. 2022), as well as embryonic development (Tani et al. 2020). Notably, R-loop depletion altered early lineage decisions, with increased expression of TFs such as *Pou3f1* and *T* in epiblast and caudal neuroectoderm, along with rewiring of the *T* gene regulatory network. At later stages, we observed elevated retinoic acid (RA) signaling from mesoderm to ectoderm, likely reinforcing ectodermal commitment. RA-related genes such as *Crabp2* and *Dlg2* were upregulated, and GRN analysis revealed expansion of *Rarb* modules in spinal cord cells. In contrast, cardiogenic networks driven by *Hand2* and *Mef2c* were diminished, suggesting impaired mesodermal potential. These findings highlight how R-loops influence not only chromatin architecture but also downstream functions such as transcriptional networks and intercellular signaling pathways that coordinate lineage specification.

Together, our results demonstrate that R-loops are critical for establishing chromatin states that prime developmental genes for activation and for maintaining balanced lineage trajectories during early differentiation. While dispensable for self-renewal, R-loops are essential for robust multilineage potential. Future studies using lineage-specific R-loop depletion or in vivo models will be important to dissect context-dependent roles of R-loops in development.

## METHODS

### Cell Culture

ZX-1 cells (Iacovino et al. 2011) (gift from Michael Kyba) were maintained in standard ESC medium (high-glucose DMEM, 10% FBS, Glutamine, MEM NEAA, β-mercaptoethanol, and LIF) on feeder cells. For feeder-free culture, cells were gradually adapted to 2i/LIF medium (NDiff 227 with LIF, 2% FBS, 3 µM CHIR-99021, 1 µM PD0325901) on gelatin-coated dishes. ZX-1 and RHKI cells were passaged every other day using accutase and plated at 2–2.5 × 10⁵ cells per 6-cm dish at 37°C, 5% CO₂. Details of RHKI cell line generation and validation are provided in Supplemental Methods.

### Gastruloid Differentiation

RHKI cells were differentiated into gastruloids as previously described (Baillie-Johnson et al. 2015; Cermola et al. 2021), with minor modifications. Briefly, cells were seeded at 300 cells/well in NDiff 277 medium in U-bottom 96-well plates. CHIR-99021 (3 µM) was added on Day 2, and medium was refreshed on Days 3 and 4. For Rnaseh1 induction, 0.5 µg/ml doxycycline was added from Day –2 to Day 5. Samples were collected at Days 0, 3, and 5 for RNA-seq and scRNA-seq. Full protocol details are provided in Supplemental Methods.

### Protein Extraction and Western Blotting

Cell pellets were lysed in buffer containing 50 mM Tris-HCl (pH 7.5), 250 mM NaCl, 3 mM EDTA, 3 mM EGTA, 1% Triton X-100, 0.5% NP-40, and 10% glycerol supplemented with 1× Halt protease inhibitor (ThermoFisher). After a 15-minute incubation on ice, lysates were centrifuged at 15,000 g for 15 minutes at 4°C. The supernatant was used for protein quantification (Bio-Rad), followed by boiling in protein loading dye prior to SDS-PAGE and Western blot analysis. Primary antibodies were used at the following dilutions: anti-tubulin (1:5000) and anti-HA (1:1000); secondary antibodies, goat anti-rabbit HRP were diluted 1:5000.

### RNA Extraction and RNA-seq

Total RNA was extracted using the Direct-zol RNA Microprep kit (Zymo) with on-column DNase digestion. Bulk RNA-seq libraries were prepared using the Illumina stranded mRNA prep kit and sequenced on a NovaSeq (PE150). For scRNA-seq, gastruloids were dissociated with accutase, and single-cell suspensions were processed using the 10x Genomics Chromium Next GEM Single Cell 3’ kit. Libraries were sequenced on a NovaSeq targeting 20,000 reads per cell. Details of bulk and sc-RNA-seq sample processing are provided in Supplemental Methods.

### DRIP-qPCR

DNA:RNA immunoprecipitation (DRIP) was performed as previously described (Sanz and Chédin 2019; Dowling et al. 2024) with minor modifications. Genomic DNA from RHKI cells (±Dox) was extracted, digested with restriction enzymes, and treated with RNase III. DNA was immunoprecipitated using the S9.6 antibody, and enrichment was quantified by qPCR.

Oligonucleotides used in this study are included in Supplemental Table S4. Full protocol details are provided in Supplemental Methods.

### CUT&RUN and MapR

CUT&RUN and MapR were performed following established protocols (Yan and Sarma 2020; Hainer and Fazzio 2019) using anti-CTCF and RHΔ-MNase, respectively. Briefly, cells were bound to Concanavalin A (ConA) beads, incubated with primary antibodies or fusion proteins, and subjected to targeted chromatin digestion. Libraries were prepared using a modified NEBNext Ultra II protocol and sequenced on a NovaSeq (PE150). Detailed procedures, including digestion conditions, library prep modifications, and sequencing parameters, are provided in Supplemental Methods.

### CUT&Tag

CUT&Tag was performed as described previously (Kaya-Okur et al. 2019), using antibodies against H3K4me3, H3K27me3, H3K27ac, H3K36me3, H4ac, H3.3, and phospho-Rpb1(Ser2). Briefly, cells were bound to ConA beads, incubated with primary and secondary antibodies, and treated with pA-Tn5 transposase. Libraries were prepared using Nextera primers and sequenced on a NovaSeq (PE150). Full protocol details are provided in Supplemental Methods.

### ATAC-seq

ATAC-seq was performed following established protocols (Buenrostro et al. 2015; Corces et al. 2017). Cells were lysed, tagmented with Tn5, and DNA was purified and amplified. Libraries were gel-purified, QC’d, and sequenced (NovaSeq, PE150; 10–20 million reads per library). See Supplemental Methods for full details.

### MNase-seq

MNase-seq was conducted as previously described (Henikoff et al. 2011). Cells were fixed, nuclei were isolated and digested with varying MNase concentrations, and DNA was purified. Libraries were gel-purified and sequenced (NovaSeq, PE150; 50–60 million reads per library). Detailed procedures are included in Supplemental Methods.

### Data Analysis

#### Chromatin Profiling Data Analysis

CUT&RUN, CUT&Tag, MapR, ATAC-seq, and MNase-seq data were processed using standard pipelines. Reads were trimmed with Trimmomatic (Bolger et al. 2014), aligned to the mm10 genome using Bowtie2 (Langmead and Salzberg 2012), and filtered for quality and duplicates. BigWig files were generated with deepTools (Ramírez et al. 2016), and peak calling was performed using HOMER (Heinz et al. 2010). Differential peaks were identified using HOMER and annotated with bedtools (Quinlan and Hall 2010) and clusterProfiler (Wu et al. 2021a). For MNase-seq, reads were stratified by fragment size, and merged BAMs were used to assess nucleosome occupancy. Chromatin states were inferred using chromHMM (Ernst and Kellis 2012), and visualizations were generated with deepTools, HOMER, and ggplot2.

#### Bulk RNA-seq Analysis

Reads were trimmed with Trimmomatic and aligned to mm10 (with annotation; Gencode vM25) using STAR (Dobin et al. 2012). Gene-level quantification was performed with RSEM (Li and Dewey 2011), and differential expression was assessed using DESeq2 (Soneson et al. 2016). Genes with adjusted p-value ≤ 0.05 and |log₂FC| ≥ 0.58 were considered differentially expressed. PCA, heatmaps, and violin plots were generated in R. Gene ontology enrichment was performed using clusterProfiler. Time-course expression trends were analyzed using DEGreport.

#### scRNA-seq Analysis

Raw reads were aligned to the mm10 genome (mm10-2020) using Cell Ranger (10x Genomics), and gene count matrices were processed using Seurat and Scanpy (Hao et al. 2024, 2021; Wolf et al. 2018). Doublets were identified with DoubletFinder (McGinnis et al. 2019), and cells were filtered based on mitochondrial content, feature count, and cell cycle scores. PCA were performed with using Harmony for batch effect correction (Korsunsky et al. 2019), followed by UMAP reduction for visualization and clustering with PARC from pyVIA (Stassen et al. 2021). Cell types were annotated using CellMatch and validated against published gastruloid and embryonic datasets. Trajectory inference was performed using pyVIA, integrating real timepoint information from our dataset and two published references.

Pseudotime was initially inferred at the cluster level using pyVIA, achieving ≥70% correlation with real timepoints. Single-cell pseudotime was then refined using Margaret (Pandey and Zafar 2022), leveraging the same adjacency matrix and timepoint alignment strategy. For comparisons between control and Dox-treated samples, pseudotime distributions and trajectory weights were analyzed without time-series constraints to avoid the bias due to limited timepoints in our dataset. Differential gene expression was assessed using MAST (Finak et al. 2015) with subsampling to control for cell number differences. Cell–cell communication was inferred using CellChat (Jin et al. 2021, 2025) with CellChatDB.mouse as the collection of ligand-receptor pairs.

Gene regulatory networks (GRNs) were inferred using a multi-step approach combining SCENIC and scMTNI, which incorporated trajectory information during GRN inference (Zhang et al. 2023; Aibar et al. 2017). Baseline GRNs were constructed from control data (ours and published), stratified by cell type, and refined through repeated subsampling. Final GRNs for control and Dox-treated conditions were derived from robust TF-target edges conserved across subsamples. Functional differences were assessed via overlap and GO enrichment analyses.

## ACKNOWLEDGEMENTS

We thank Michael Kyba for ZX-1 cells. We thank I.-H. Wang for advice on analysis of sc-RNA-seq data. We thank J. Benanti, A. Herchenröther, S. Gopalan, E. Hass, T. Basak, and P. Gautam for critical comments on the manuscript. This work was funded by grants R01HD072122 and R01HD104971 from the NIH. C.-H.C. helped conceive of the study, designed experiments, performed all experiments, performed all data analyses, prepared all figures, and helped draft the manuscript. T.G.F. acquired funding, helped conceive, design, and oversee the study and helped draft the manuscript.

## SUPPLEMENTAL FIGURES

**Supplemental Figure S1.**
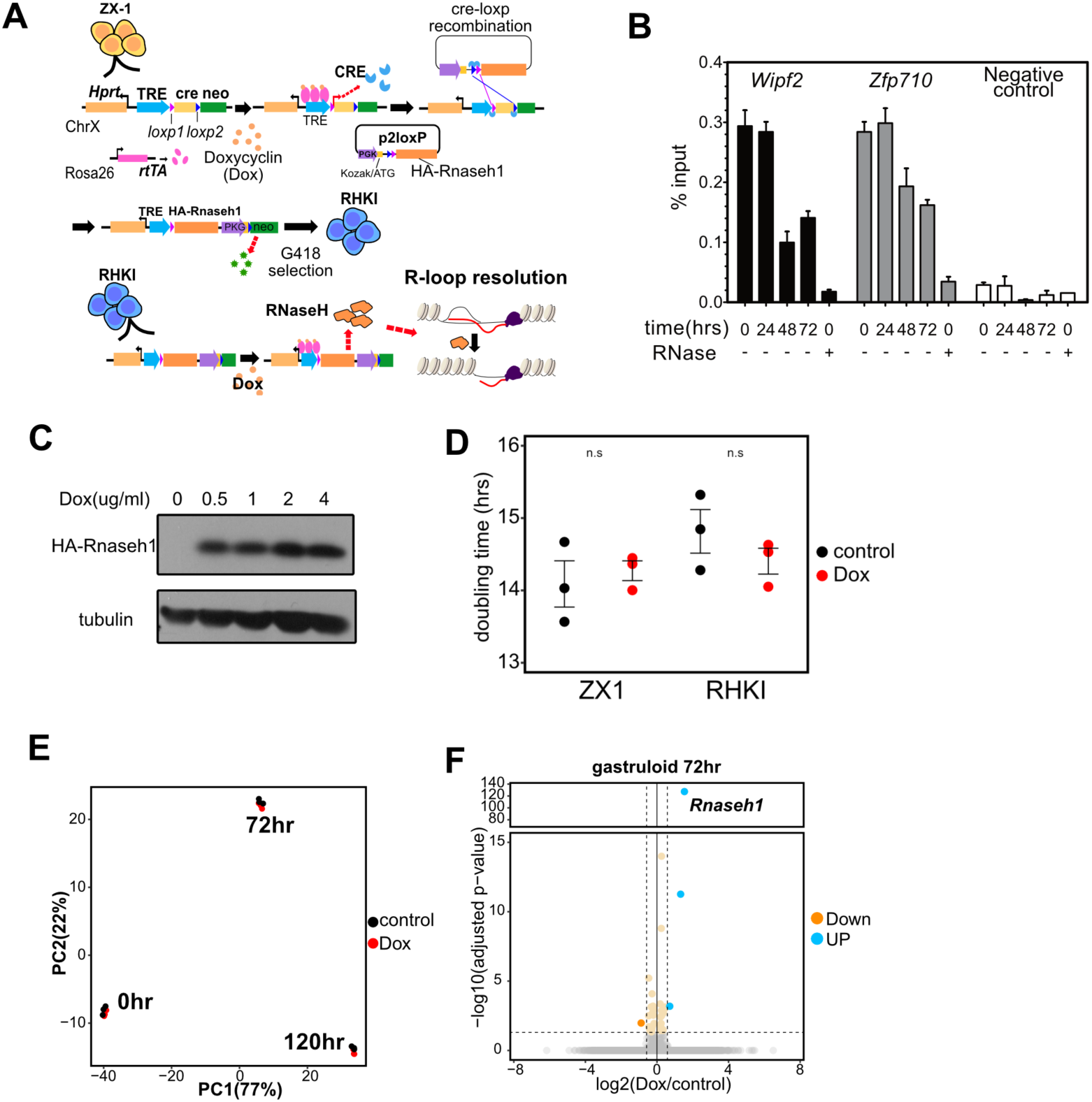
Effects of R-loop disruption on mESCs during self-renewal and differentiation. (A) Schematic for generation of inducible RNaseh1 line (RHKI). The RNaseh1 expression cassette is knocked in upstream of the *Hprt* locus in parental (ZX-1) mESCs, removing the Cre cassette and allowing ectopic expression of HA-*RNaseh1* by doxycycline. See Experimental Procedures for details. (B) Timecourse of R-loop depletion in response to Dox treatment in RHKIs. Shown are DRIP-qPCR measurements for indicated loci. “RNase” depicts a control in which gDNA was treated with RNaseA and RNaseH before DRIP. Shown are the mean (percent of input in IP) and standard error of mean. (C) Western blot of HA-RNaseH1 showing induction at different concentrations of Dox. Tubulin is shown as a loading control. (D) The doubling time of ZX-1 and RHKI between control and Dox condition. Shown are the mean and standard error of mean for each. Three replicates were performed for each condition. (E) PCA plot of bulk-RNA-seq data from three timepoints post gastruloid differentiation in RHKIs. (F) Volcano plot of bulk RNA-seq for 72 hour gastruloids with the cutoff: |log2FC| ≥ 0.58 and adjusted p-value ≤ 0.05.

**Supplemental Figure S2.**
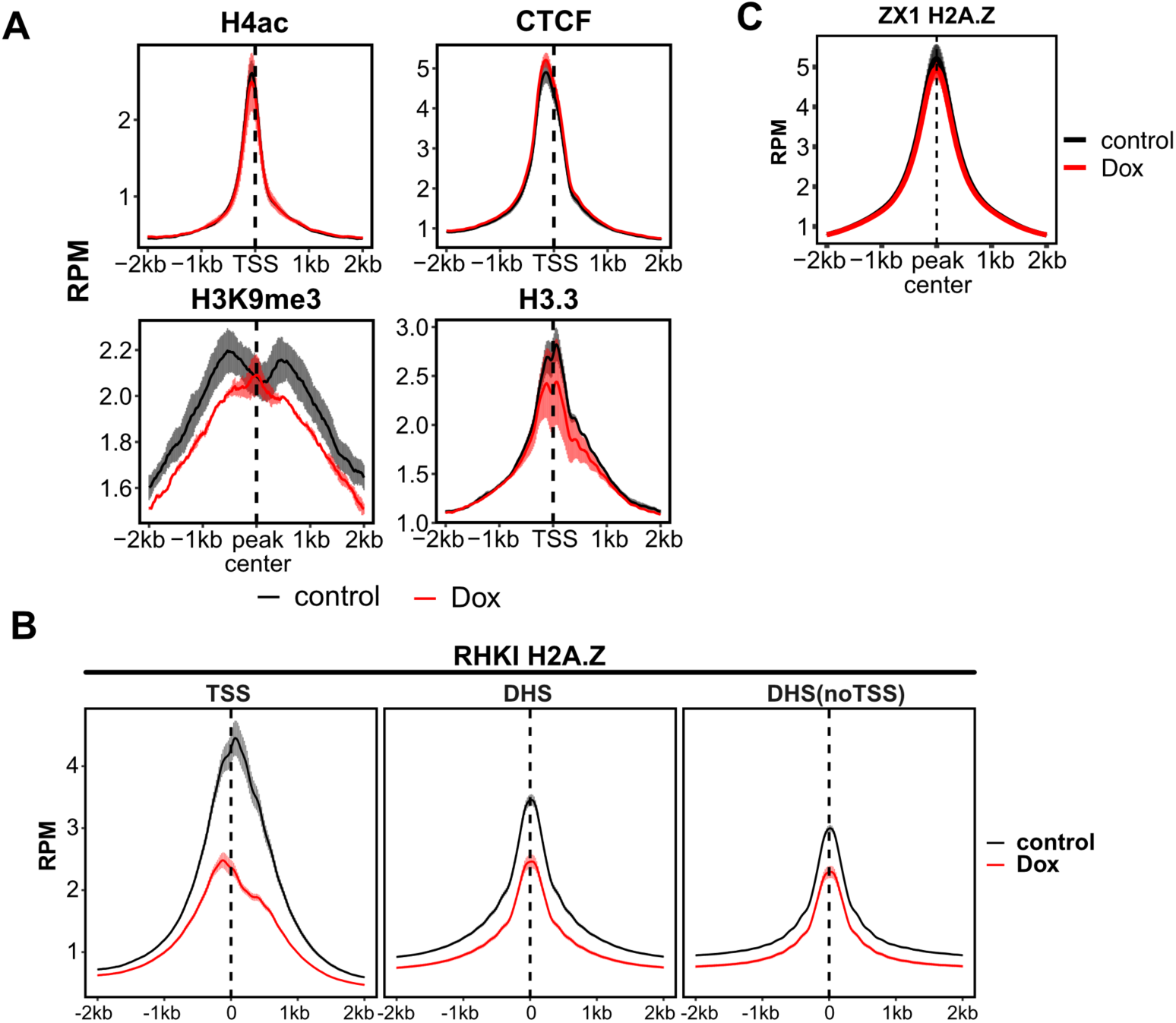
Effects of R-loop depletion on epigenomic features. (A) Aggregation plots of epigenetic features across TSSs (H4ac, CTCF, and H3.3) and peak centers (H3K9me3: GSE32218) between control and Dox cells. Shown are mean RPM and standard error of mean. (B) H2A.Z enrichment across TSSs, DHSs (w/ and w/o TSSs) between control and Dox conditions. Shown are mean RPM and standard error of mean. (C) Aggregation plot of H2A.Z enrichment in parental (ZX-1) cells, comparing control and Dox conditions. Shown are mean RPM and standard error of mean.

**Supplemental Figure S3.**
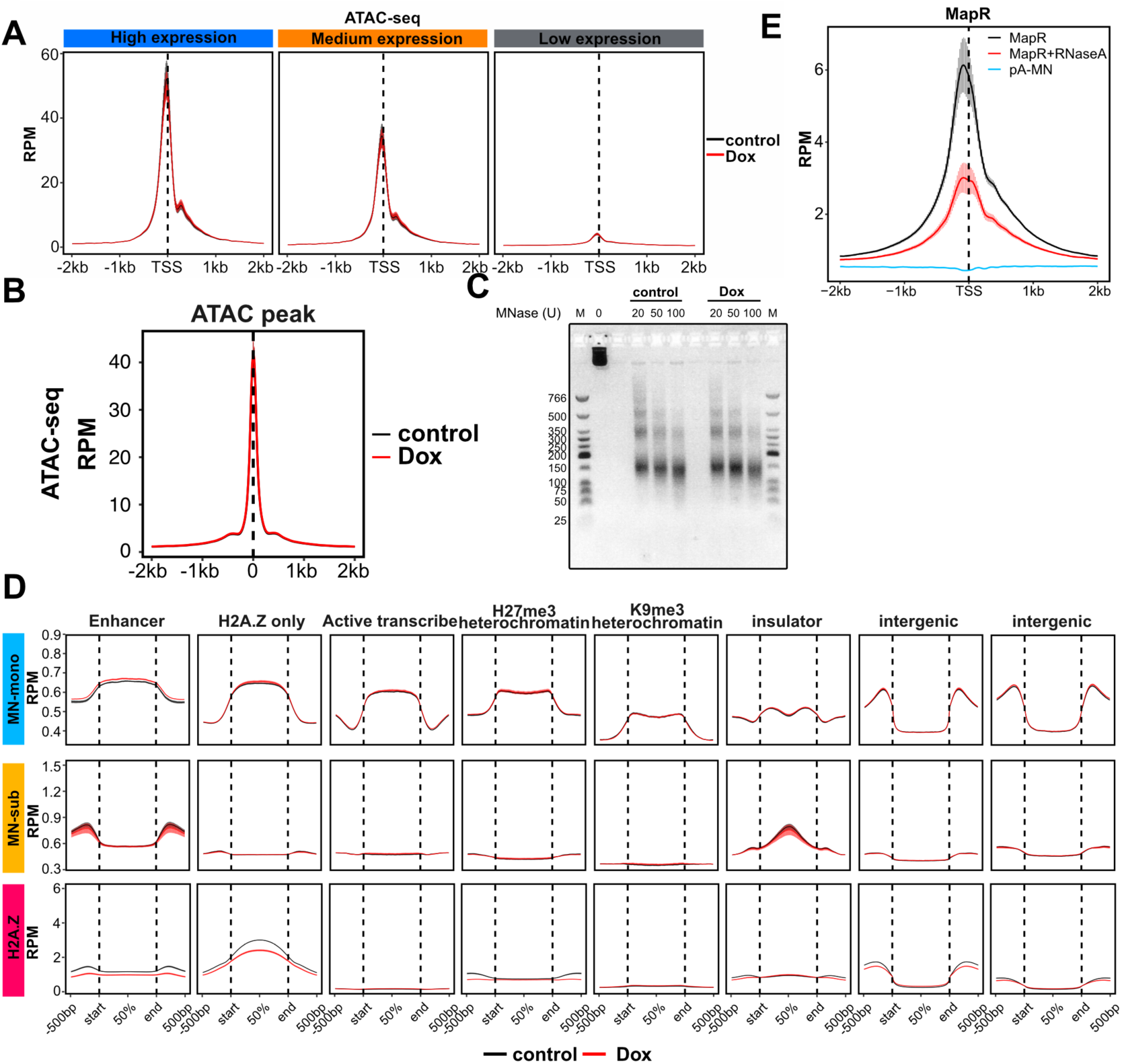
Changes in ATAC-seq and MNase-seq profiles upon R-loop depletion. (A) Aggregation plots of ATAC-seq signal (1-120bp fragment size) across TSSs split into high, medium, and low expression categories as in Figure 3B. Shown are mean RPM and standard error of mean. (B) Aggregation plot of ATAC-seq signal (1-120bp fragment size) across ATAC-seq peaks. Shown are mean RPM and standard error of mean. (C) Size distribution of DNA fragments from MNase titration for MNase-seq experiment. 20U, 50U, 100U of MNase were used for both conditions. M: low molecular weight ladder. 0: undigested control. (D) Aggregation plot of the remaining eight states from ChromHMM that were not shown in Fig. 3C, showing enrichment of MN-mono, MN-sub and H2A.Z for each state. Shown are mean RPM and standard error of mean. (E) Aggregation plot of MapR (R-loop mapping) across TSSs in RHKIs and is shown with mean and standard error of mean. Exogenous RNaseA treatment and pA-MN (without RNA-DNA hybrid binding domain) are as negative controls.

**Supplemental Figure S4.**
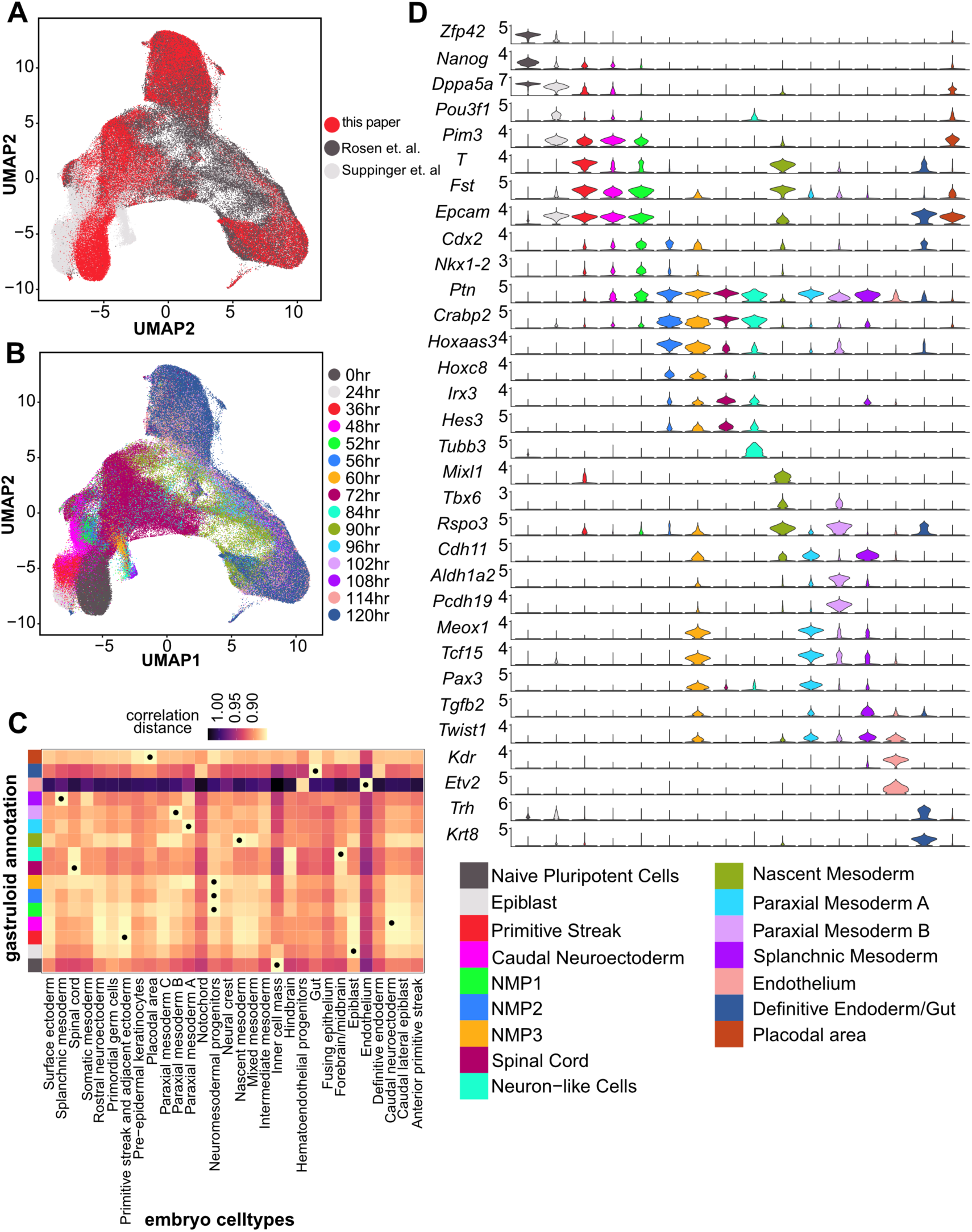
Cell annotations and marker genes for each cluster. (A) and (B) UMAP plot for the combination of two published datasets and our data. (A) is labeled by data source and (B) is labeled by time point. (C) Heatmap showing annotation results by comparison with scRNA-seq data from mouse embryos (E3.5 to E8.5). The color code for each cell type in gastruloids is shown in (D). Dots indicate the most correlated cell type between gastruloids and embryos (the lower of the value, the closer the cell types). (D) Violin plot of marker genes across all cell types in the combined scRNA-seq data.

**Supplemental Figure S5.**
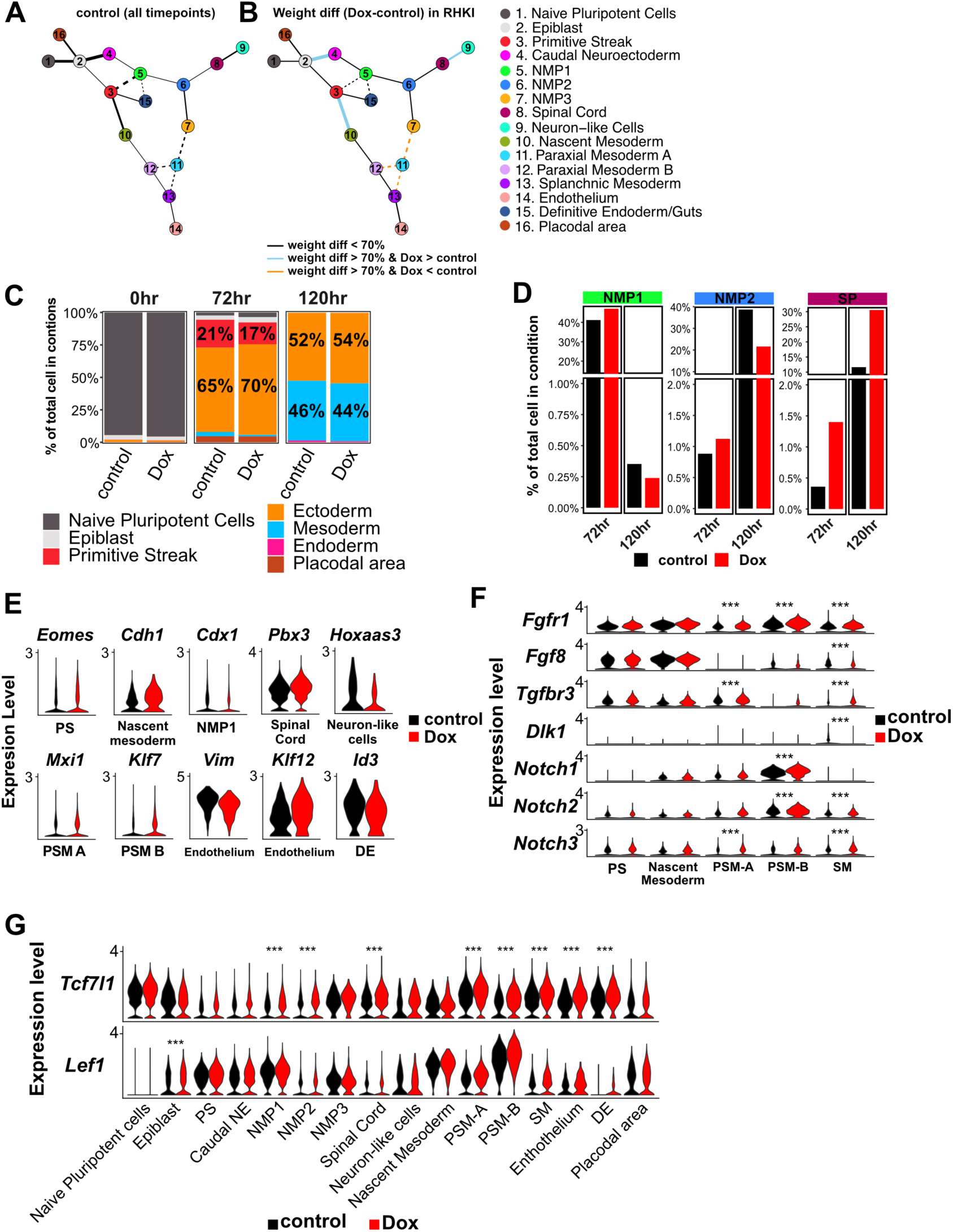
Changes in lineage trajectory, cell proportion, and gene expression resulting from R-loop depletion. (A) Comprehensive lineage trajectory plots from all control data, combining gene expression similarity and real timepoint/pseudotime information (see Experimental Procedures). Each node is labeled as in (B). The thickness/weight of edges indicate the likelihood of connection between nodes. Edges with dashed lines indicate other possible routes that are not in the final trajectory shown in Figure 5C. (B) Difference in lineage trajectory between control and *Rnaseh1* overexpressing cells. Each edge represents the weight difference between control and Dox and indicates the changes in likelihood between the nodes when comparing control and Dox. A difference of less than 70 quantiles (see Experimental Procedures) is indicated in black. If the difference is larger than 70 quantiles, Dox > control is depicted as blue and Dox < control is orange. (C) The lineage proportion changes between control and Dox in RHKI at each timepoint during gastruloid differentiation. Ectoderm: Caudal Neuroectoderm, NMP1, NMP2, NMP3, Spinal Cord and Neuron-like cells. Mesoderm: Nascent Mesoderm, Paraxial Mesoderm A/B, Splanchnic Mesoderm and Endothelium. Endoderm: Definitive Endoderm/Guts. (D) The cell proportion between control and Dox in NMP1, NMP2 and Spinal Cord at 72 hours and 120 hours in RHKIs. (E) Violin plots of DEGs in different cell types with the same criteria in Figure 5E. (F) Violin plots of gene expression related to signaling pathway in the mesoderm trajectory. *** indicates adjusted p-value < 0.0001. (G) Violin plots of WNT signaling effectors, *Tcf7l1* and *Lef1*, across all cell types in RHKI upon gastruloid differentiation. *** indicates adjusted p-value < 0.0001.

**Supplemental Figure S6.**
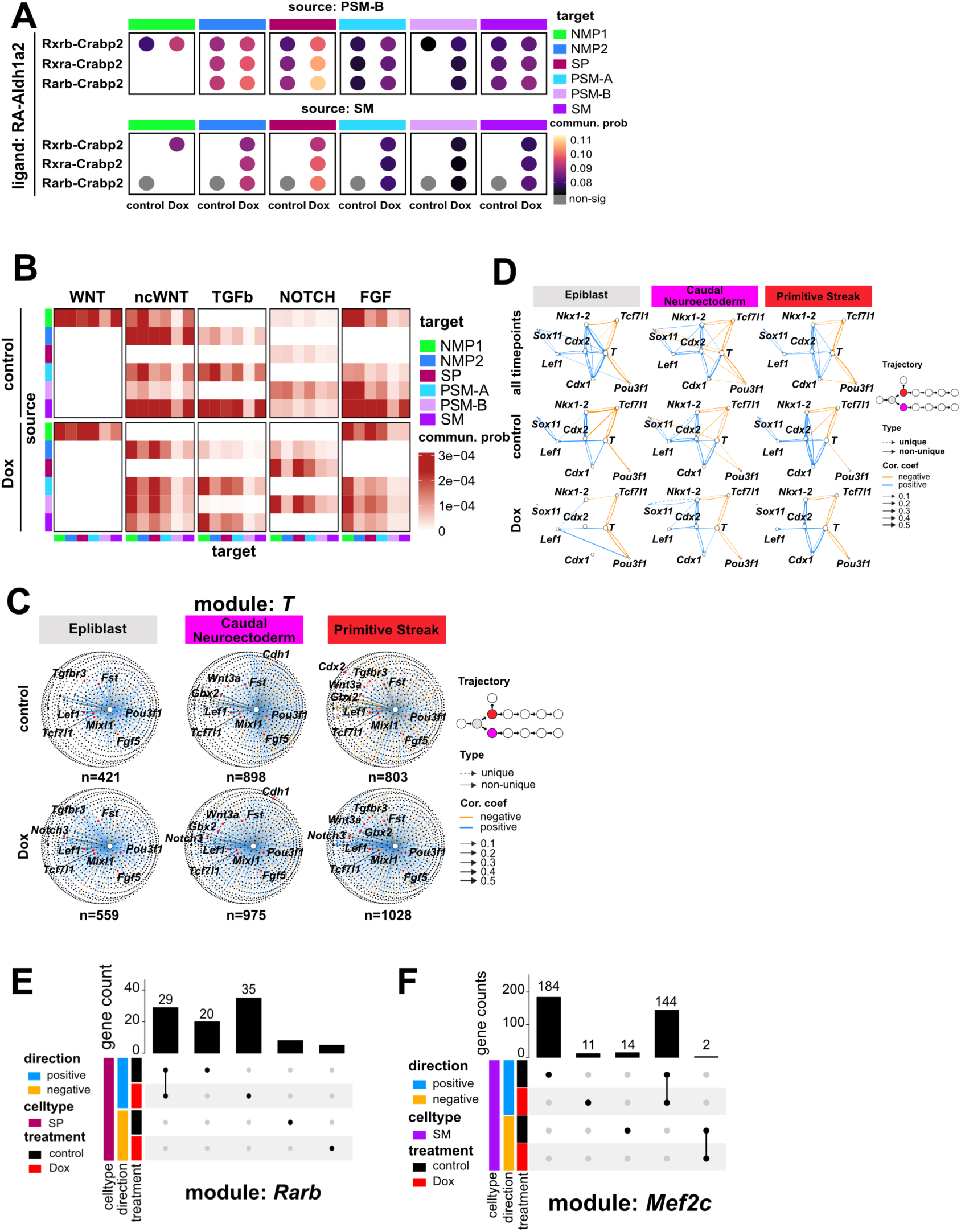
Alterations in CCC and GRNs upon R-loop loss. (A) The communication probability of ligand-receptor pair in retinoic acid signaling from PSM-B and SM to both mesoderm cells (PSM-A, PSM-B, SM) and ectoderm cells (NMP1, NMP2, spinal cord). (B) The heatmap of communication probability of WNT, ncWNT, TGFβ, NOTCH and FGF signaling between control and Dox among the combination of mesoderm cells (PSM-A, PSM-B and SM) and ectoderm cells (NMP1, NMP2, SP). (C) Network plots illustrating TF-target relationships in the *T* module. The edge type, color, and weight are the same indication as shown in Figure 6D. The trajectory diagram highlights the position of the cell type in trajectory map especially here the early fate decision point of gastruloids. (D) The correlation network plot among critical TFs in celltypes at early stage. The arrowhead of the edge indicates the directionality of TF → target relationship. All control indicates our control dataset and two published datasets for network analysis. The edge type, color and weight are the same indication as shown in Figure 6D. (E) and (F) Upset plots shows the intersect targets among different category, *Rarb* module in (E) and *Mef2c* module in (F). Direction indicates the positive or negative correlation between TF and targets.

## SUPPLEMENTAL METHODS

### RHKI Cell Generation

Feeder-free ZX-1 cells were seeded at 1–1.5 × 10⁵ cells/well in 6-well plates with 2i/LIF medium and 1 µg/ml doxycycline (Dox) for CRE induction. Cells were transfected the next day with p2loxp-HA-Rnaseh1 (constructed by cloning a synthetic HA-tagged Rnaseh1 cDNA lacking the mitochondrial targeting sequence into p2loxp-EGFP at the EcoRI site) using FuGENE HD in OptiMEM. After 24 hours, cells were serially diluted and selected with 300 µg/ml G418. Colonies were expanded and screened by Western blot for Dox-inducible HA-Rnaseh1 expression. Clones were also assessed for ESC morphology and pluripotency marker expression by qPCR.

### Gastruloid Differentiation (Full Protocol)

Cells were seeded at 4 × 10⁴ per 6-cm dish on Day –4. On Day –2, medium was replaced with fresh 2i/LIF. On Day 0, cells were detached with accutase, washed with PBS, and seeded at 300 cells/well in 40 µl NDiff 277 medium in untreated U-bottom 96-well plates. On Day 2, 150 µl NDiff 227 medium with 3 µM CHIR-99021 was added. Medium was refreshed on Days 3 and 4. Doxycycline (0.5 µg/ml) was added from Day –2 to Day 5. NDiff 227 was thawed overnight at 4°C and pre-warmed before use.

### Bulk RNA-seq Library Preparation

1 µg RNA per sample was used with the Illumina stranded mRNA prep kit (10 PCR cycles). Libraries were QC’d via Bioanalyzer and Qubit HS, and sequenced on a NovaSeq (PE150), yielding 20–30 million reads/sample.

### scRNA-seq Library Preparation

Gastruloids were dissociated with accutase (5–10 min at 37°C), washed with PBS, and resuspended in 0.04% BSA/PBS. Viability was assessed with trypan blue. Cells were loaded into the 10x Genomics Chromium system targeting 10,000 cells/sample. Libraries were prepared per manufacturer’s protocol, QC’d, and sequenced on a NovaSeq (PE150), targeting 20,000 reads/cell.

### DRIP-qPCR

RHKI cells were lysed in buffer (10 mM Tris, pH 7.5; 10 mM EDTA; 10 mM NaCl; 0.5% sarcosyl) with Proteinase K at 37°C overnight. DNA was extracted via PCI and ethanol precipitation, then digested with BsrGI, EcoRI, HindIII, SspI, and XbaI, followed by RNase III treatment. DNA was split into aliquots with or without RNase H (and RNase A if needed), re-extracted, and immunoprecipitated with 10 µg S9.6 antibody in MeDP buffer overnight at 4°C. Complexes were pulled down with Protein G beads, washed, and eluted in STOP buffer at 65°C. DNA was purified and analyzed by qPCR using SYBR Fast reagent and specific primers. Enrichment was calculated as % input and fold change over negative control.

### CUT&RUN and MapR

RHΔ-MNase was purified as described(Yan and Sarma 2020). Cells (2 × 10⁵) were washed and bound to activated ConA beads, blocked, and incubated overnight at 4°C with anti-CTCF (1:100) or RHΔ-MNase in antibody buffer. For CUT&RUN, cells were incubated with pA-MNase, washed, and resuspended in dig-wash buffer. For MapR, RHΔ-MNase-bound cells were washed and resuspended similarly. Digestion was initiated with 3 µl 100 mM CaCl₂ on ice for 30 minutes. Reactions were stopped with STOP buffer and incubated at 37°C. DNA was purified via PCI and ethanol precipitation. Libraries were prepared using NEBNext Ultra II with TruSeq adapters (no USER), purified with Ampure XP beads (0.9×), amplified (14 cycles), gel-extracted (150–650 bp), QC’d, and sequenced (NovaSeq, PE150; 5–10 million reads/library). Antibodies used in CUT&RUN and the adapter/primer sequence for library prep (TruSeq) are listed in Supplementary Table S4.

### CUT&Tag

2 × 10⁵ cells were bound to ConA beads and incubated overnight at 4°C with primary antibodies (1:50) in antibody buffer (wash buffer + 0.05% digitonin, 0.1% BSA, 2 mM EDTA). The next day, Guinea Pig anti-Rabbit IgG (1:100) was added for 1 hour. After washing, pA-Tn5 (1:50) was added in dig300 buffer and incubated for 1.5 hours. Tagmentation was performed in dig300 buffer with 10 mM MgCl_2_ at 37°C for 1 hour. Reactions were stopped with TAPS/SDS buffers, followed by quenching and PCR amplification (17 cycles). Libraries were purified with Ampure XP beads (1.1×), QC’d, and sequenced (NovaSeq, PE150; 5–10 million reads/library). Antibodies used in CUT&Tag and the adapter/primer sequence for library prep (Nextera) are listed in Supplementary Table S4.

### ATAC-seq

1 × 10⁵ cells were lysed in buffer (10 mM Tris-HCl, pH 7.5; 10 mM NaCl; 3 mM MgCl₂; 0.1% NP-40; 0.1% Tween-20; 0.01% digitonin) on ice for 15 minutes. After washing, cells were tagmented in TD buffer with Tn5 at 37°C for 30 minutes. DNA was purified (MinElute), amplified (8–10 cycles), gel-purified (150–500 bp), QC’d, and sequenced (NovaSeq, PE150; 10–20 million reads/library).

### MNase-seq

1 × 10⁶ cells were fixed with 1% formaldehyde for 15 minutes, quenched with glycine, and nuclei were isolated in NE buffer. Nuclei were digested with MNase (10–100 U) for 5 minutes at 37°C. Reactions were stopped with STOP buffer, and DNA was purified as in CUT&RUN. One microgram of DNA was used for library prep, gel-purified, QC’d, and sequenced (NovaSeq, PE150; 50–60 million reads/library).

